# Endorsing Darwin – Global biogeography of the epipelagic goose barnacles *Lepas* spp. (Cirripedia, Lepadomorpha) proves cryptic speciation

**DOI:** 10.1101/019802

**Authors:** Philipp H. Schiffer, Hans-Georg Herbig

## Abstract

**Aim:** We studied different species of gooseneck barnacles from the globally distributed rafting genus *Lepas* to examine whether the most widespread species are true cosmopolitans and to explore the factors influencing the phylogeny and biogeography of these epipelagic rafters.

**Location:** Temperate and tropical parts of the Atlantic, Pacific, and Indic oceans.

**Methods:** We used a phylogenetic approach based on mitochondrial 16S and coI sequences, and the nuclear 18S gene to elucidate patterns of inter- and intra-species divergence. Altogether, five species of *Lepas* from 18 confined regions of the Atlantic, Pacific and Indic oceans were analyzed.

**Results:** A combination of nuclear and mitochondrial sequences provided robust phylogenetic signals for biogeographic classification of subgroups in *Lepas* species. *Lepas australis*, restricted to cold-temperate waters of the southern hemisphere shows two separate populations in the southern hemisphere (coastal Chile, other circum-Antarctic sampling sites) most probably related to temperature differences in the southern Pacific current systems. A more complex differentiation is seen for the cosmopolitan *L. anatifera* that thrives in warmer waters. In total, it is differentiated in four regional subgroups (coastal Chile, Oregon, Indopacific in general, Atlantic) and a global group, which might be either an ancient stemgroup, but more probably is an anthropogenic artefact. Separation into subgroups likely reflect geological vicariance effects. For *Lepas*, these were closure of the Isthmus of Panama, installation of the cool Benguela Current in the later Miocene including its persistence into the Present, Pleistocene current systems in the western Pacific differing from today, and in general lowered Pleistocene temperatures, and finally present-day current systems. The extreme ecological generalists *L. anserifera* and *L. pectinata* are not differentated according to available data and might represent true global species. Data for *L. testudinata* remain ambiguous.

**Main conclusions:** Our data indicate cryptic speciation in some but not all species of the cosmopolitan epipelagic genus *Lepas.* Regional distribution of genetically different populations rely on a wealth of in part interacting inherited geological factors and modern traits. Controlling factors differ between species according to their ecospace limits and demonstrate the need for studies on species level to avoid unjustified generalizations. Data indicate that allopatric speciation is the main mode of divergence. True global species, if existent at all, need to be extreme ecological generalists, like *Lepas pectinata* and *L. anserifera*.

## Introduction

Before starting his work on the ‘Origin of Species’ Charles Darwin spent several years studying barnacles (Cirripedia, Thoracica). He published four monographs on these cosmopolitan, highly derived, sessile crustaceans (Darwin, 1851; 1854). In the volume dedicated to lepadomorph (or goose) barnacles Darwin (1851) noted that ‘*all the valves, even in the same species, are subject to considerable variation in shape*’ and, ‘*from the foregoing description it will be seen how extremely variable almost every part of this species is’*. This later led others to suspect either cryptic speciation or great morphological plasticity in at least the type species *Lepas anatifera* (Newman & Ross, 1971; Newman, 1972). In fact, the cosmopolitan epipelagic rafter *Lepas* might constitute a model genus to test speciation patterns in the open ocean and to verify the existence of truly cosmopolitan species.

Since beginning of the 21st century complexes of cryptic species have been discovered in a growing number of taxa by the application of the Phylogenetic Species Concept. While the most popular showcase might have been the African Elephant (Roca et al., 2001; 2005), the marine environment habours a large and diverse number of taxa for which cryptic species have been identified. In a now classic paper Knowlton (2000) reviewed cryptic speciation in the marine environment and discussed the consequences on our understanding of the nature and age of species boundaries. She, and later Westheide & Schmidt (2003), addressed the fact that neglect of abundant cryptic speciation leads to serious underestimation of global diversity and thus has impact on present taxonomic usage and on the question of the existence of cosmopolitan species (Norris, 2000). On the other hand Coyne & Orr (2004) stated that ‘*[…] it is not clear how barriers to gene flow operate in the open ocean’*, and therefore the oceans of the world might constitute an interconnected environment where gene flow between populations might persist. Consequently, a first question arises, whether the sea might habour global/cosmopolitan species with little or no genetic differentiation (Hewitt, 2000), or if divergence into geographically distinct cryptic species is common in most marine taxa - despite apparent morphological uniformity in some. A subsequent second question tackles the problem of speciation mechanisms, i.e. speciation by sympatry in an apparently barrierless environment versus speciation by allopatry in an environment with hitherto unrecognized barriers (see discussion in Norris & Hull, 2011).

Especially taxa spending their whole life or very long times in the open ocean – such as holoplanktonic organisms, pelagic swimmers, drifters and rafters, and organisms with extended pelagic larval stages – might maintain gene flow over enormous distances in the oceanic environment. These taxa might have the potential to be truly cosmopolitan species, but if no effective gene flow is maintained, diversification and evolution of regional variants should be expected. In that context, the matter of time is controversely discussed. Indeed it has been proposed that rapid evolution into regional sub-types should be common in the zooplankton (Peijnenburg & Goetze, 2013), but vice versa, Norris & Hull (2011) stressed the importance of time for speciation. Thus, the key to our understanding of speciation and species boundaries in the open oceans lies within these organisms.

Surprisingly, the majority of studies up to date deal with geographically restricted benthic species, most with planktonic larvae. These organisms, however, have to return to a suitable benthic environment within the finite larval lifetime and must comply with the requirements of their - often very complex - lifecycles and are thus not ideal candidates to resolve the question of cosmopolitanism. Among the many examples available, we mention the ascidian *Ciona intestinalis* (Caputi et al., 2007), the shallow-water temperate to boreal bryozoan *Electra pilosa* (Nikulina et al., 2007), the pantropical seaweed *Halimeda* (Kooistra et al., 2002; Verbruggen et al., 2005) and the pantropical sea urchins *Eucidaris* and *Tripneustes* (Lessios et al., 1999; 2003), which all show separation by vicariance induced by land barriers and/or ocean currents.

Among the analysed pelagic organisms, the hydrozoan *Obelia geniculata* showed previously unrecognized speciation between oceanic regions. It is, however, absent from some major oceanic regions (Govindarajan et al., 2005) and, therefore is not a suitable taxon for a global assessment. A global phylogeographic study of the bryozoan *Membranipora membranacea* commonly attached to rafting substrate substrata (Schwaninger, 2008) indicated a high degree of regionalism in the ocean induced either by temporal vicariance, i.e. allopatric speciation, due to climatic events or by founder populations dispersing to a new suitable habitat. Copepods are intensively studied and yielded differing results. Goetze (2003) found high levels of cryptic diversity at the species level in the Eucalanidae and concerning *Rhincalanus nasutus* geographically distinct populations within the Pacific and between the Pacific and the Atlantic. Using the coI gene as a single marker, Goetze (2005) found that in two antitropical *Eucalanus* species differentiation might be due to habitat discontinuities induced by large oceanic gyre systems. Eberl et al. (2007) showed genetic diversification in the ‘*pseudobenthic*’ rafting copepod *M. gracilis,* but could not resolve the question of cryptic speciation, as the study is based on a single mitochondrial locus. Population differentations within and between oceans were later also described in other copepod species (Goetze, 2011; Norton & Goetze, 2013; Halbert et al., 2013; Cornils & Held, 2014; Andrews et al., 2014). Yet, such an entrapment in ocean current systems was questioned by Provan et al. (2009) concerning the copepod *Calanus finmarchicus* based on genetic data and modelling of palaeodistribution during the Last Glacial Maximum. Also data on the planktonic nudibranch genus *Glaucus* yielded different results on the species level: the Indo-Pacific *G. marginatus* shows four regional populations, whereas the cosmopolitan *G. atlanticus* appeared to be panmictic (Churchill et al., 2013). However, refined analysis of *G. atlanticus* showed the separation of two Indopacific and two Atlantic haplotypes, related to the gyres of the northern and southern hemispheres.

Concerning protists, extensive work on planktonic foraminifer genera show cryptic ongoing speciation and regionalism from genetic and fossil data on a global and local scale (Darling et al., 1999; de Vargas et al., 1999; 2002; Darling & Wade, 2008; Aurahs et al., 2009; Weiner et al., 2014). Results for the planktonic diatom *Pseudo-nitzschia pungens* show limited gene flow and strong isolation by distance (Casteleyn et al., 2010). Protists, however, might be more strongly affected by temperature, salinity, chemistry and other ecological factors and/or higher speciation rates and thus results might not be analogous metazoans.

All presently available data stress the fact that the planktonic environment is much more complex than previously anticipated (see also Darling & Wade, 2008). Most studies find consistent differences between the major ocean basins (e.g. Norton & Goetze, 2013), but high levels of connectivity have been reported for the Antarctic ocean (Bortolotto et al., 2011).

Reports about putative panmictic species in the epipelagic environment are not only scarce but, beyond that, studies to date do not show a coherent picture of speciation on a global level in the epipelagic environment. In consequence, more research is needed to understand the mechanisms that either permit cosmopolitan distribution or lead to cryptic speciation in the epipelagic environment, as already emphasized e. g. by Saez & Lozano (2005).

The goose barnacles of the Crustacean genus *Lepas* (Cirripedia, Thoracica; Fig. 1) are ideal candidates to investigate the question of cosmopolitan species and speciation in the sea since they are epipelagic surface drifters with abundant populations inhabiting all oceans (Fig. 1). As mentioned above, reasons to suspect cryptic speciation in the genus go back to Darwin’s descriptions and later to Newman & Ross (1971) suggesting sub-species inhabiting different oceanic regions. Here we test the possible divergence of these epipelagic rafters on a global scale. We used geographically most distant sampling sites, i.e. two from the opposite sites of the North Atlantic and two from the southwestern and southeastern Pacific, as well as several additional sites across the globe to address the question of cryptic speciation or true cosmopolitanism in independent species of one genus. These are the putative cosmopolitans *L. anatifera*, *L. anserifera* and *L. pectinata*, as well as *L. australis* inhabiting the circum-subantarctic seas and the rare *L. testudinata*.

**Figure 1:**
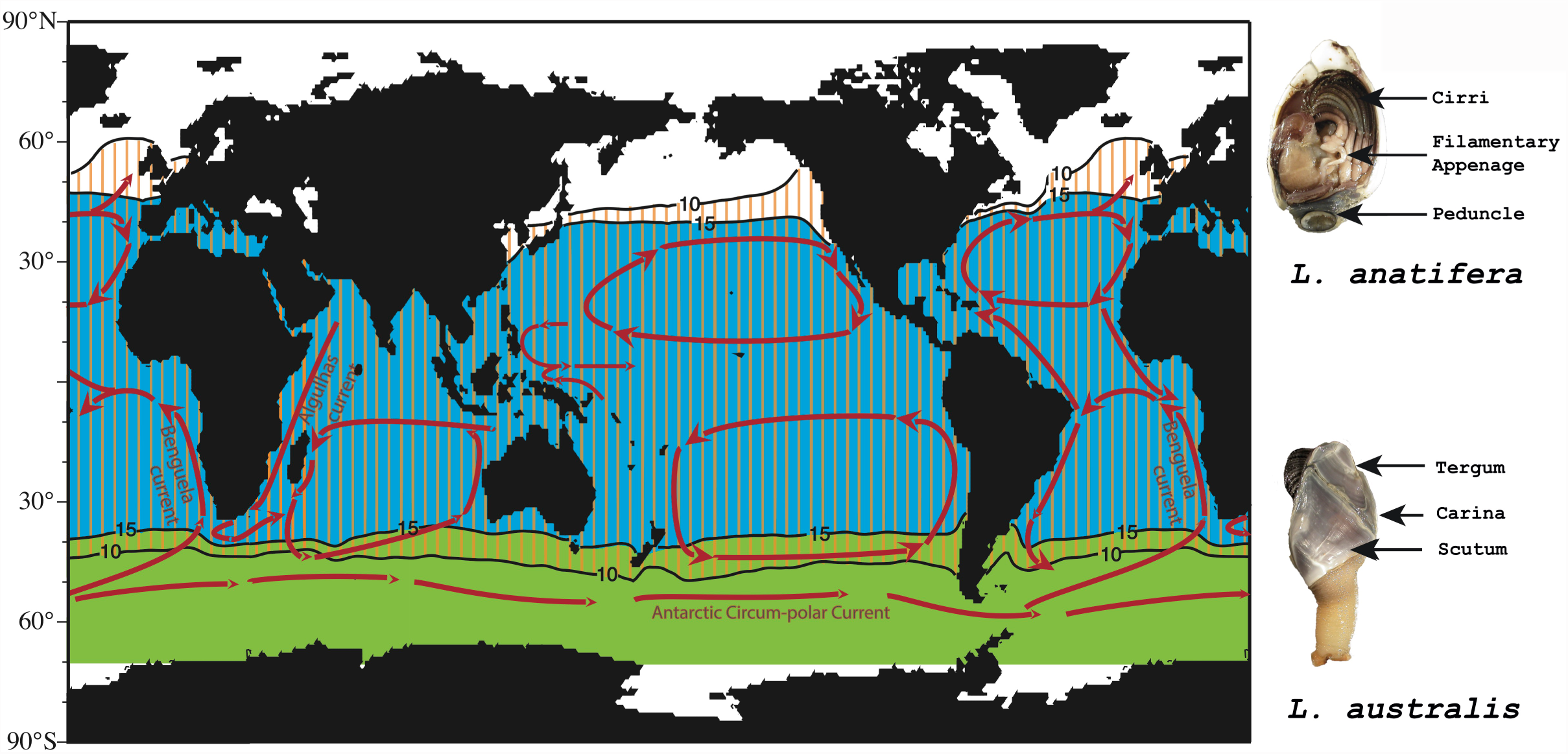
Major oceanic current systems and the distribution of *Lepas anatifera* (blue), *L. australis* (green), and *L. pectinata* (orange, dashed) after (Hinojosa et al., 2006). L. *anatifera* appears restricted to waters warmer than 15°C, while *L. australis* is found in cooler water masses.

We show clear phylogenetic diversification, i.e. cryptic speciation, in two species, but also indications for gene flow between major oceans. The extended geographic sampling sites allow discussion of pathways or possible barriers to gene flow, i.e. the acting mechanism of speciation, in the open ocean. We find that allopatric speciation (due to vicariance by temporal introduction of barriers or dispersal: oceanic currents, passive rafting on driftwood, icebergs, kelp, etc.) is the most likely explanation for divergence in different *Lepas* species, but cannot rule out sympatric speciation as a possibility in the pelagic environment.

## Materials and Methods

### Sampling and initial Species Identification

Samples were collected from around the globe (see Table 1) between summer 2007 and summer 2012. Specimens were preserved in Ethanol (EtOH) immediately after collection. EtOH was exchanged upon sample reception and the material was then stored at −20°C. Individual specimens were again placed into EtOH and stored at −80°C after DNA extraction. All original species identifications from local collectors were (re-)confirmed by the first author in accordance with descriptions from the literature (Darwin, 1851; Gruvel, 1905; Nilsson-Cantell, 1930; Newman, 1972; Anderson, 1994; Hinojosa et al., 2006). Diagnostic features like the form and number of filamentary appendages, as well as shell structures (e.g. number/presence/location of umbonal teeth, form of shell plates, ridges and imprintings on these plates) were analysed with the aid of stereo-microscopes and photo-captured for a subset of the samples.

**Table 1:** Areas from first sampling campaign. Numbers are given for specimens for which all 3 loci could be sequenced, total number of specimens analysed is thus higher.

| Region | Specific location | *L. anatifera* | *L. anserifera* | *L. australis* | *L. pectinata* |
| --- | --- | --- | --- | --- | --- |
| Coastal Chile | Coquimbo | 16 |  | 4 | 8 |
| Gulf of Mexico | Florida Coast | 10 | 5 |  |  |
| Spain | Ria de Vigo | 16 | 1 |  | 1 |
| New Zealand | N/A |  |  | 6 | 6 |

### DNA Extraction and Chemistry

Muscle tissue for DNA extraction was either taken from the peduncle or the base of the cirri. DNA was extracted using commercially available kits. DNA was stored at −20°C during the experimental phase and later transferred to −80°C. Polymerase chain reactions (PCR) were conducted under the inclusion of high fidelity polymerases. When necessary 10% Trehalose or Betaine were added to enhance PCR yield and specificity. 16S and 18S Primers as described in Simon-Blecher et al. (2007) were used. A new, slightly degenerated, forward primer (FwcoInt_Lep: 5'-ATAYTAATTCGTGCWGAACTMGG) had to be designed for the coI locus while the ‘universal’ reverse primer (Hebert, 2003) could be employed. PCR set-ups and cycle settings were optimized for each marker and depending on the quality of DNA extractions. PCR products were visualised on Agarose gels and products showing neither multiple bands nor excessive smearing on the gel were cleaned employing the Exo/Sap protocol Werle et al. (1994), or commercial kits from various manufacturers. In few cases were multiple bands persisted after adaptation of the PCR protocols bands were gel-excised and purified with commercial kits. Sequences were either obtained through in-house Sanger sequencing and read out on ABI capillary DNA Analyzers at the Cologne Center for Genomics (CCG, University of Cologne), or by sending dried PCR products to Macrogen Inc. (Seoul, South Korea and Amsterdam, The Netherlands). Sequencing was in general conducted bi-directional and a subset of samples was repeatedly amplified and sequenced to control for possible sequencing errors. For 16S and coI the same forward and reverse primers as used in the PCR were employed. For the longer stretch of the 18S gene internal primers as described in Simon-Blecher et al. (2007) were used. An additional primer (18S_Fsq4_Lep: 5’-ATCGACTGGAGGGCAAGC), yielding a larger region of internal overlap, was designed.

### Genetic Markers and Outgroup

We constructed a three loci phylogeny, sequencing fragments of one nuclear and two mitochondrial genes. We used the canonical barcoding gene cytochrome c oxidase subunit I (coI), which had successfully been applied in analyses of cryptic species complexes (Hebert et al., 2004). The 16S ribosomal DNA (16S rDNA or 16S) gene, had been employed in recent studies on barnacle phylogeny (Simon-Blecher et al., 2007). To avoid depicting only the maternal site of evolution, we added a fragment of the conserved 18S rDNA (18S rDNA, SSU) nuclear gene that previously had been used in establishing a barnacle phylogeny (Simon-Blecher et al., 2007). From our initial analysis of the full (∼1,800bp) sequence of the 18S gene we were able to define a smaller region that contained enough informative sites to robustly delimit species and subsequently amplified this shorter fragment. We obtained specimens of *Dosima fascicularis* from Argentina and *Heteralepas* sp. from Madeira as outgroup species for the phylogenetic analyses.

### Phylogenetic Analysis

Sequences were compared with barnacle sequences stored at the NCBI nucleotide database with BLAST to detect possible contamination with foreign DNA. All sequences obtained were submitted to *GENBANK* and may be retrieved under the accession numbers: FJ906772 - to FJ906777 and GU993588 to GU993704, as well as XXXXXX. Acquired sequence traces were visually inspected with the programs Codon Code Aligner (v.2.0.6) and Geneious versions 5 and 6. In the Codon Code Aligner the Phred (Ewing et al., 1998) quality-assessment algorithm was applied to control the chromatograms. Assemblies were then built under application of the phrap sub-program and visually re-inspected for ambiguous base calls, which were then corrected, or else the sequence discarded. For Geneious we used stand alone Phred if necessary and then assembled the reads with the implemented algorithm. We initially re-constructed a phylogeny for a limited set of samples available to us at that time using maximum likelihood and Bayesian approaches as described in the Supplementary Methods. After acquiring our full set of samples we combined these with the initial alignments of each single locus in Geneious and re-aligned with clustalΩ 1.0 (Sievers et al., 2011). We visually inspected and edited the alignments where necessary in Geneious and SeaView v.4 (Galtier et al., 1996). The Geneious software was also used to create a combined all loci alignment. We used SplitsTree v.4 (Huson, 2008) on the combined alignmet to explore phylogenetic signal in our data and assess potential conflicts between loci and sample groups. We then choose to analyse all loci individually. We ran PhyML (release 20131016) (Guindon & Gascuel, 2003) on our data choosing the GTR + gamma model and used RAxML in version 7.7.2 (Stamatakis, 2006) running for 20 iterations as second validation on some alignments. Bootstrapping was conducted using for 1000 generations in PhyML. The program was also used to re-confirm the inference of our initial geographically restricted data set originally performed in PAUP*. We tested our alignments with modelgenerator v.0.85 (Keane et al., 2006) and implemented the evolutionary models in MrBayes 3.2.2 (Huelsenbeck & Ronquist, 2005) to infer Bayesian phylogenies. We ran this program for 5 Million generations in 4 parallel runs on the local computer cluster CHEOPS. For *L. anatifera* population we constructed haplotype genealogies using Fitchi (Matschiner, 2015), which also provided us with Fst values for haplotypes in the different oceans.

### Results

All *Lepas pectinata* and *L. australis* specimens shared morphological features as described in literature and could therefore assigned to the respective species. *L. australis* specimens from Coquimbo in Chile were much smaller than other individuals of this species and therefore considered as juveniles. *L. testudinata* specimens were identified by A. Biccard (Cape Town). Specimens of *Lepas anatifera* from the Southeast Pacific were juveniles as well, thus differing in size from specimens collected from other regions. Interestingly, all *L. anatifera* from the Gulf of Mexico, as well as some specimens from other regions show at least one line of brownish squares across the scuta (and sometimes across the terga as well) and slightly more elevated radial lines (or ribs) on the scuta (see Fig. 1 for shell nomenclature). Both these features were not present in the same extent in the remaining samples. Nevertheless, all samples were clearly assigned to the species *L. anatifera*, as the varying morphological features were previously described (see especially Newman, 1972).

### Initial phylogenetic analysis – indications from the Atlantic and Southern Pacific

In a first assay we analysed specimens from three oceanic regions (Schiffer & Herbig, 2008), the North Atlantic (including Gulf of Mexico), the Southeast Pacific (Coquimbo, Chile) and and the Southwest Pacific (New Zealand). From these sampling sites genomic DNA from 76 specimens was suitable to sequence all three loci (Table 1, Supplementary Excel File 1). Across all species out of the 76 specimens analysed, 74 differing mitochondrial haplotypes for both marker loci (16S and coI) were obtained, while only six 18S haplotypes were retrieved for the whole sample set. The length of the alignment of all loci was 2,792 bp. Maximum likelihood (ML) and Bayesian inference for the all-loci alignment yielded the same tree topology on genus, species and intra-species level. Our later re-calculation of the ML tree with PhyML showed no deviation in topology from the first inferences. Only minor differences are present in the placing of single individuals within biogeographical subgroups between ML and Bayesian inference. Monophyly of *L. pectinata*, *L. anatifera* and *L. anserifera* received 100% bootstrap and posterior probability support, while *L. australis* is supported with 99.5 and 100%, respectively (Fig. 2).

**Figure 2:**
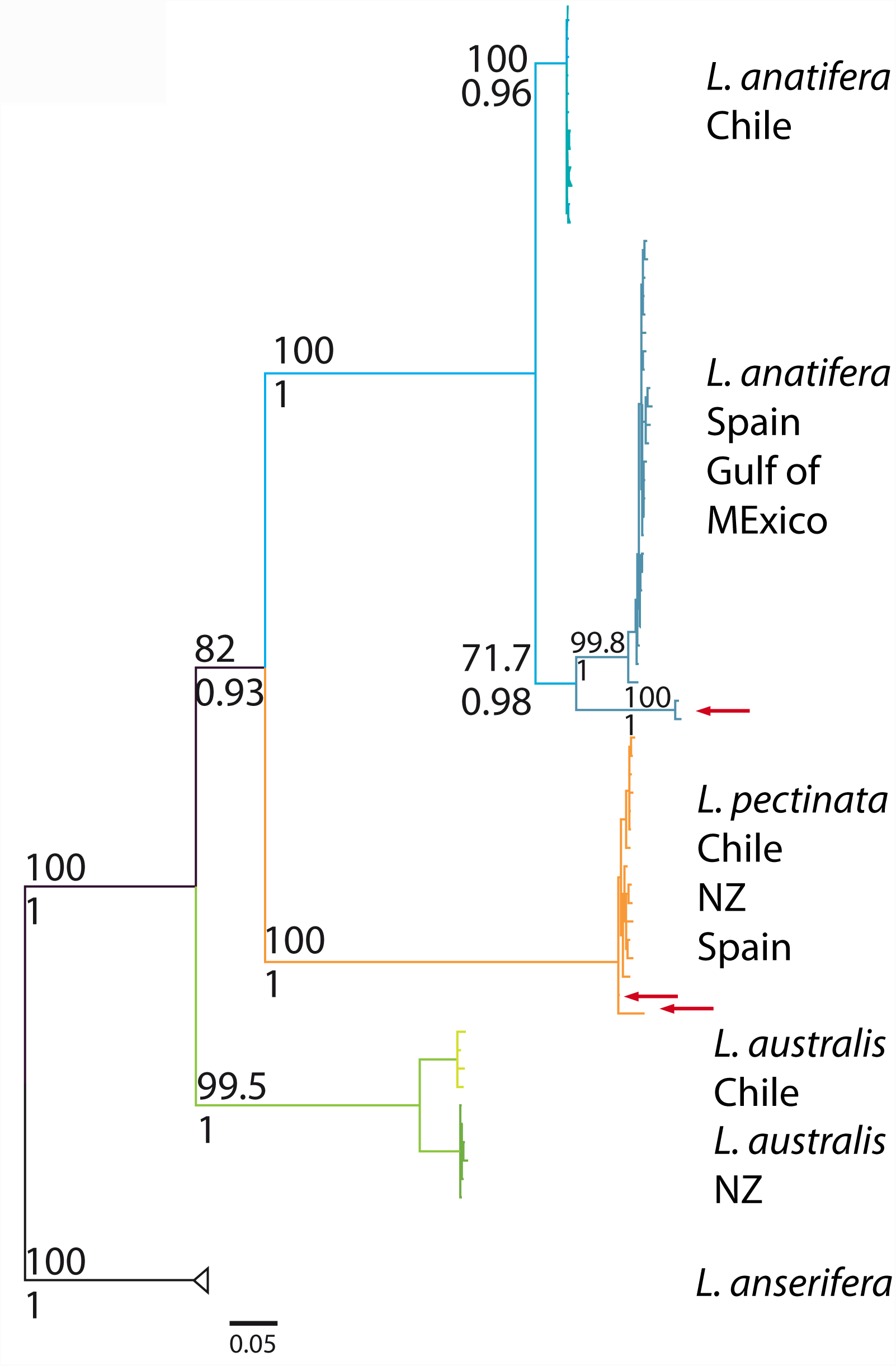
Initial phylogeny based on three genes (18S, 16S, coI) from a subset of samples from major oceanic regions re-constructed using PAUP* (re-confirmed with PhyML) and MrBayes. A split in sub-types is found in *L. anatifera* and *L. australis.* The red arrows indicate “outliers” in *L. anatifera* and *L. pectinata*. In *L. anatifera* these were later found to belong to a global group, while in *L. pectinata* no obvious biogeographic subgroup was found.

On an intra-species level a split was found between the *L. anatifera* populations from the Southeast Pacific (monophyletic with 100% bootstrap support and 96% posterior probability) and populations from the Northeast Atlantic/Gulf of Mexico (71.7 and 98%) (Fig. 2). A group of specimens from the Gulf of Mexico (both 100%) was set apart from the remaining individuals of the Northeast Atlantic/Gulf of Mexico clade (99.8 and 100%). One *L. pectinata* specimen from the Northeast Atlantic and Southwest Pacific, was set apart from all other individuals of this taxon (Fig. 2), albeit only weakly supported (76 and 50%). Two distinct biogeographical monophyletic clusters were found in *L. australis,* where specimens from the SE Pacific form one (99.9% and 99%) and from the SW Pacific (99.9 and 100%) a second group (Fig. 2).

Refined phylogenetic analysis – towards a global biogeography

### Nuclear phylogeny

To map the diversity indicated by our first analysis more finely on a global scale and to resolve population structures we acquired samples from all major oceanic regions (Figure 3, Table 2). As in the first assay, most specimens could be collected for *L. anatifera* and least for *L. pectinata* (Supplementary File 1).

**Figure 3:**
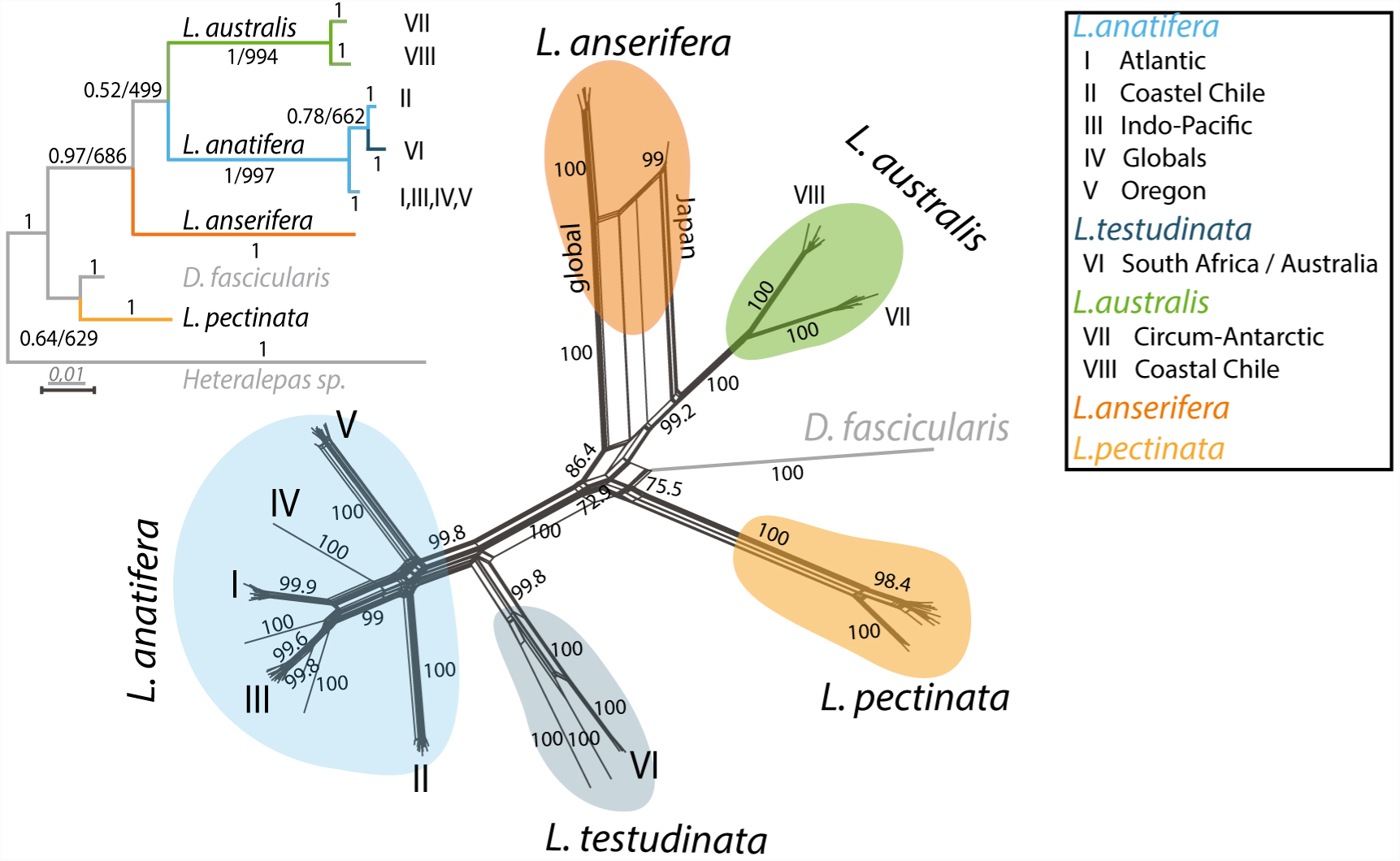
Phylogenetic (ML, Bayesian) tree based on analysis of a fragment of the 18S ribosomal gene including two outgroup species. Posterior probabilities and bootstrap values are printed to the edges, where support is maximal only the former are given. A Neighbournet constructed in SplitsTrees indicates a clear treelike signal in our sequence data, but also conflicts in the *L. anserifera* data, which led us to analyse mitochondrial loci independently.

**Table 2:** Major oceanic regions and sampling points per species from the second sampling campaign.

| Oceanic region | Coastal location | *L. anatifera* | *L. anserifera* | *L. australis* | *L. pectinata* | *L. testudinata* |
| --- | --- | --- | --- | --- | --- | --- |
| “Chile” | Coquimbo, Chile | + |  |  | + |  |
| South Pacific | Chile, south/off-coast | + |  | + |  |  |
|  | New Zealand |  | + | + |  |  |
| North East Pacific | Oregon | + |  |  |  |  |
| North West Pacific | Japan |  | + |  |  |  |
| Equatorial Pacific | Tonga | + |  |  |  |  |
| East Indian Ocean | Western Australia | + |  | + |  | + |
| South Africa | South Africa | + | + | + | + | + |
| Gulf of Mexico | Florida Coast | + | + |  | + |  |
| Atlantic | Argentina |  |  | + |  |  |
|  | Uruguay |  | + |  |  |  |
|  | Brasil | + | + |  |  |  |
|  | Senegal |  | + |  |  |  |
|  | Cape Verde | + | + |  |  |  |
|  | Madeira | + |  |  |  |  |
|  | Spain | + | + |  | + |  |

From the initial phylogenetic analyses we were able to define a shorter region of the 18S gene that carried enough signal for species and possibly sub-species discrimination. We acquired 18S sequences from 140 specimens additional to those used in the first analysis that were suitable to be incorporated into our biogeographic analysis. In this set we found two 18S variants in *L. anatifera*, two in *L. australis*, and one each in *L. pectinata*, *L. anserifera*, and the outgroups. Specimens morphologically identified as *L. testudinata* were very similar in the sequence of this gene to the *L. anatifera* specimens. The phylogenetic analysis of these sequences yielded a species tree (Fig. 3), which reconfirmed our initial findings from fewer sampling sites (see above), but includes the outgroups *D. fascicularis* and *Heteralepas* sp. The tree shows a split into geographical subgroups in *L. anatifera* and *L. australis*, while no such pattern is retrieved in the other species. Posterior probabilities strongly support placing and monophyly of all species, but indicate that the groupings of *L. australis* and *L. anatifera* as well as *L. pectinata* and *D. fascicularis* as sister species cannot be resolved with this locus alone. As expected when using a shorter and highly conserved gene sequence, bootstrap values drop in comparison to what was described above for our initial phylogenetic inference. However, *L. australis* and *L. anatifera* still receive very strong support and the other species are far from being weakly supported (Fig. 3). The inferred grouping is also in line with a recent revision of barnacle phylogeny analysing the evolution of parasitic species (Rees et al., 2014).

In *L. australis* we found two groups, one encompassing specimens sampled from the Chilean coast alone, while the second group contains samples from New Zealand (sampled at different times), Argentina and originating of the western coast of Australia. Finally, we find a single sample from South Africa in this group. The single group inferred for *L. pectinata* contains specimens from New Zealand and Chile (both from different sampling campaigns), as well as the Western Mediterranean (Baleares, Ibiza) and the Azores. The group found for *L. anserifera* holds samples from the North Atlantic (Gulf of Mexico, Northeast Spain, Senegal, the Cape Verde Islands), South Atlantic (Brasil, Uruguay, South Africa), and New Caledonia, western Australia, and Japan.

Using the 18S tree as an indication of possible intra-species divergence and a phylogenetic backbone we tested our mitochondrial data for potential conflict in the phylogenetic signal. A neighbournet constructed from the all-loci alignment in SplitsTree (Fig. 3) shows a tree-like structure in most of its branches. However, some potential conflicts were indicated (e.g. for *L. anserifera* in a Japanese sample and in the *L. testudinata* subgroup). Further it appeared likely that the coI and 16S loci evolve at different speeds. Therefore we decided to infer individual phylogenies for each of the two mitochondrial loci.

### Mitochondrial phylogenies

The coI and 16S trees support all *Lepas* and outgroup species with high posterior probabilities and bootstrap support, but differ in their positioning of single species (Figure 4). In particular in the Bayesian analyses the outgroup *Dosima fascicularis* (from the coast of Uruguay) is positioned next to *L. australis* and *L. anserifera* in the 16S tree, while it is found akin to *L. testudinata* and thus close to the *L. anatifera* group in the coI tree. However *L. anserifera* is also found closer to *L. anatifera* in the coI tree, where it is nested inside of an *L. pectinata* and *L. australis* split, than is seen in the 16S tree. Here *L. pectinata* is closest to *L. testudinata*, which is an outgroup to *L. anatifera*. These general groupings also slightly deviate in the maximum likelihood inference (Supplementary Figures 2 and 3).

**Figure 4:**
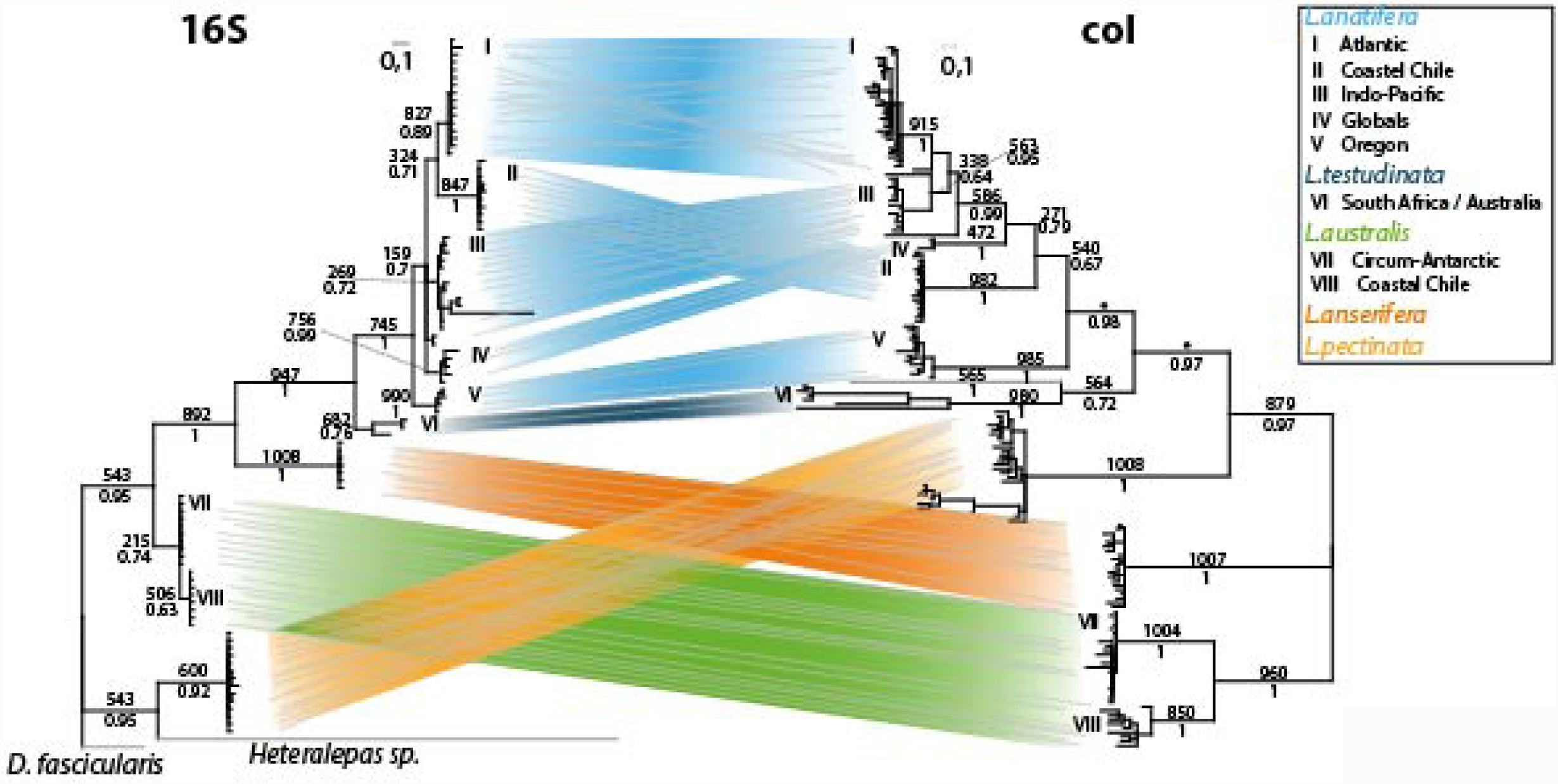
Comparsion of mitochondrial single locus phylogenies based on PhyMl and MrBayes. All major groupings are retrieved from both loci. The faster evolving coI gene provides more intra-group resolution. Bootstrapping values are given above, posterior propabilities below branches.

Both genes support a split in *L. australis* already described for the 18S gene above, one group from coastal Chile opposed to one encompassing specimens from the other circum-Antarctic sampling sites. The split however is far more pronounced in the coI gene, indicating the higher substitution rate of this locus.

The mitochondrial genes support the evolution of *L. anatifera* into several biogeographical sub-groups, but disagree in the placing of groups in relation to each other. Consequently, branches leading towards single groups have strong support, while branches separating groups have weaker posterior probabilities and bootstrap values. The Chilean subgroup, already inferred with the 18S gene, is retrieved in both mt gene trees. It is clearly separated from the other groups, but is supplemented by a single sample from eastern South Africa. Conversely, one single individual sampled at the Chilean coast plots in a group with samples from the Indo-Pacific region, namely South Africa, the Easter Islands, Tonga, Japan, and the Juan Fernandez Islands off the Chilean coast. Supposedly both non-fitting individuals were transported out of their natural habitats. Two further groups are separated from the others. These are specimens from the Oregon Coast (the North East Pacific) and a group with samples from the Atlantic Ocean. The Atlantic group had already been inferred in our initial survey (Schiffer & Herbig, 2008) and is now supplemented by specimens from Brasil, Madeira, and further samples from Spain (Cádiz). Also in our initial survey, two specimens from the Gulf of Mexico were set apart from all other *L. anatifera* sampled there and at the Spanish Atlantic coast (Fig. 1). Interestingly, these two specimens are now placed in a larger group of individuals sampled around the globe (Tonga, Cape Verde Islands, western Australian coast, and one sample from South Africa/Cape Town).

As already indicated above, both phylogenies based on these genes place *L. testudinata* as an outgroup to *L. anatifera*. This is in congruence with the inferred 18S gene tree. Inside *L. testudinata* we find a split between samples from South Africa and Australia.

In *L. pectinata* we retrieved a rather unstructured tree topology for both genes, but see at least an indication for a possible sub-group inside the globally distributed samples in the coI. This group holds samples from South Africa, the mid Atlantic regions (Azores), northeast Spain and the western Mediterranean (Ibiza).

In *L. anserifera* the coI gene seems to indicate the possibility for sub-populations, but our data is clearly not sufficient to resolve this.

To further elucidate the complex biogeographic pattern found in the mitochondrial *L. anatifera* phylogenies we calculated Fst values and a haplotype networks for the coI gene. In contrast to many other studies we did not rely on reticulation networks to resolve haplotype connectivity, we implemented a novel method to construct haplotype genealogies (Fig. 5) based on Salzburger et al. (2011) that incorporate the phylogentic results and should generate a biologically more meaningful picture (Matschiner, 2015). We found that this method reflects very well the pattern in the phylogenies but yields some additional information through the possibility of comparison inside monophyletic subgroups. In total we analysed 576 residues in the coI gene for 25 samples from the Atlantic, 10 from the Gulf of Mexico, 7 from South Africa, 19 from coastel Chile, 6 from the Equatorial Pacific, 5 from the East Indic, 3 from the South Pacific, 3 from the North East Pacific, and added 6 *L. testudinata* specimens. Pairwaise results are summarised in Supplementary Table 2. Here we found that panmixis between Atlantic and Gulf of Mexico is confirmed with very low Fst values, while the separation of the Oregon and coastal Chile groups from other regions is firmly established. It appears that gene flow persists between South Africa and the South Pacific (containing Easter Islands) and Equatorial Pacific (containing Tonga) persists, as indicated to a combined Indo-Pacific in the phylogenetic analysis (Fig. 4).

**Figure 5:**
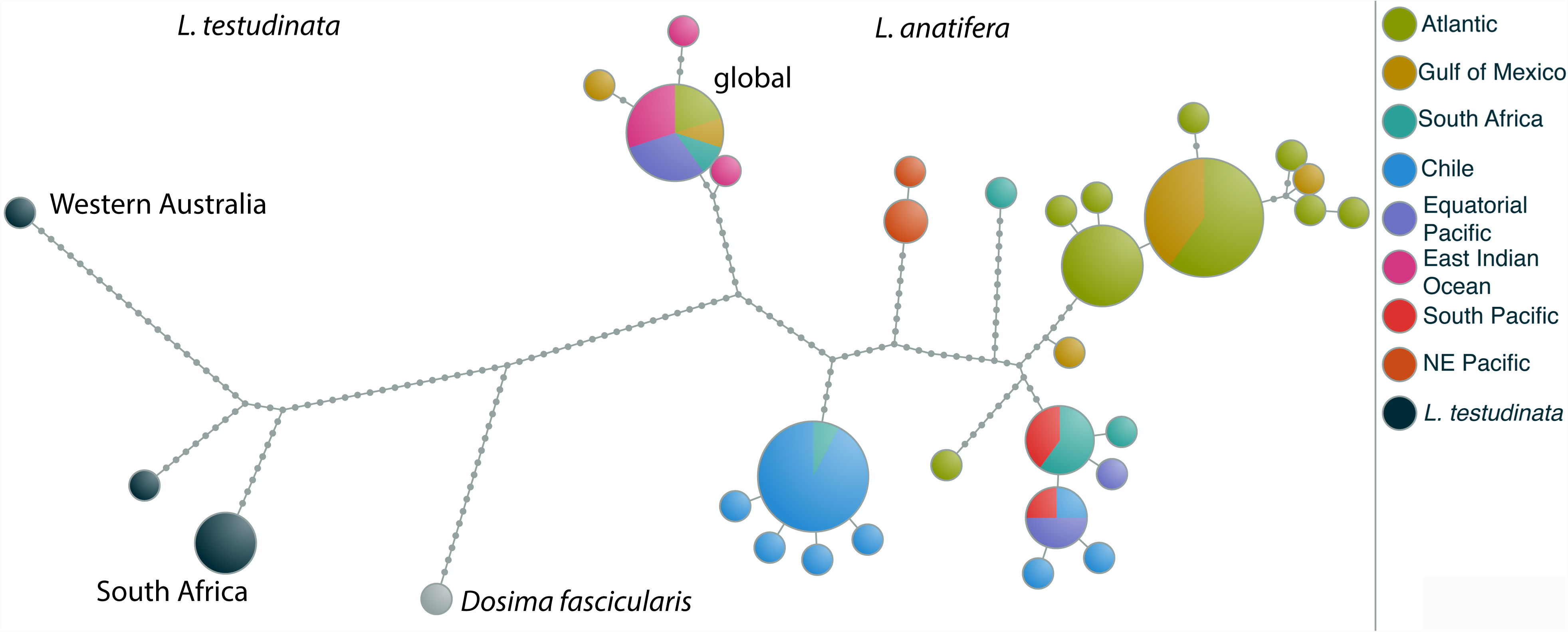
Haplotype genealogy of the coI of the *L. anatifera* and *L. testudinata* populations computed with Fitchi. A separation of oceanic regions described in the main text and depicted in Fig. 6 can be seen, as well as one “global” group in *L. anatifera*. The *L. testudinata* haplotype recovered from Australia is set apart from the South African ones.

## Discussion

### Global biogeography

Barnacles within the genus *Lepas* attach to flotsam (wood, algae, styrofoam, buoys, etc.), vessels and swimming animals after spending up to two months drifting as larvae (Anderson, 1994) (Darwin, 1851; Hinojosa et al., 2006). *Lepas anatifera* is the goose barnacle with the highest abundance worldwide (Young, 1990), as reflected in our sampling. It thrives where the sea surface temperature (SST) exceeds 18°C (Hinojosa et al., 2006) and is thought to survive in waters warmer than 15°C (Patel, 1959). Temporary settlements as transient species at the limits, resp. outside its contiguous distribution is known (Skinner & Barboza, 2014). *Lepas pectinata,* on the other hand, tolerates somewhat cooler temperatures (Zevina, Memmi, 1981). The geographic ranges of both species reach from approximately 50-55° N to 40-45° S (temperature/latitude relation after Locarnini et al., 2006 fig. 1), thus spanning across temperate and tropical regions of the world’s oceans. *Lepas australis* is a cold-water species inhabiting all oceans around the Antarctic continent (Hinojosa et al., 2006; Fraser et al., 2011). These ecological constraints strongly control the global biogeography of the species.

It has been suggested that species in the marine zooplankton are capable of rapid adaptation to locally restricted habitats in the oceans and thus the diversity of life in the open oceans is still grossly underestimated (Peijnenburg & Goetze, 2013). The entrapment of passively drifting populations in oceanic gyres (Palumbi, 1994), which persisted through extended geological periods (Talley, 2001), could be responsible for the existence of biogeographic provinces (Schopf, 1979), but if those systems are an important factor in general is still ambiguous. In fact, our results suggest different evolutionary trajectories in the *Lepas* species analysed. Both *L. pectinata* and *L. anserifera* appear to have one global genotype in the nuclear 18S gene. In both species the mitochondrial loci also do not suggest clear geographic subdivision, although there is the possibility of a distinct coI genotype in an Atlantic group of *L. pectinata*. In contrast *L. anatifera* and *L. australis* show local subgroups in each, the nuclear and mitochondrial data. In *L. australis* all markers support two groups, one with an exclusively Chilean genotype and one encompassing the remainder of samples around the Antarctic circle. The situation in *L. anatifera* is more complex based on the mitochondrial genes. The nuclear marker in *L. anatifera* supports three genotypes, one holding the *L. testudinata* subtype, one from the Chilean coast, and one with specimens from all major oceanic regions. However, mitochondrial data indicate a distinct population structure with some geographically restricted groups (Figs. 4, 5), thus similar to cryptic species in the copepod *Paracalanus parvus* (Cornils & Held, 2014), but we also find one global group.

### Lepas australis

It is most surprising to find a geographic pattern in *L. australis* that is even supported by the slowly evolving nuclear locus. The species in general is restricted to cold water masses linked to Antarctica, an area where longitudinal landmasses that could impede gene flow and thus lead to the evolution of geographically confined populations are not known. Contrary, the West Wind driven Antarctic Circumpolar Current (ACC) effectively mediates transport of organisms around Antarctica. Within the ACC the Antarctic Polar Front (APF, respectively Antarctic convergence, U.S. Geological Survey 2012) which coincides approximately with the winter sea ice edge and a sudden change in seawater surface temperature acts as an effective latitudinal barrier between water masses and also between marine organisms (Thornhill et al., 2008). Further north, the Subantarctic Front (= subtropical convergence, U.S. Geological Survey 2012), i.e. the northern boundary of the ACC is a second major latitudinal water-mass barrier. The effectiveness of these fronts as barriers to gene flow has been shown e.g. in the chaetognath *Eukrohnia hamata* (Kulagin et al., 2014).

All our sampling sites are situated north of the ACC. There, we recognize a common population of *L. australis* in the cool to temperate waters north of the Subantarctic front, termed herein the ‘Southern Ocean subgroup’. That population of *L. australis* is obviously caught in the current system adjoining north of the subtropical convergence, albeit also circling eastward around Antarctica, and in the northeast deviating currents systems, which sweep up the coasts of Western Australia and New Zealand (South Indian Current s. l.), respectively Argentina and South Africa (Benguela Current).

The ‘coastal Chile subgroup’ differs from all other sampling sites. It matches biogeographic results of a separate eastern Pacific/Chilean province from other pelagic organisms. Examples are the copepod *Rhincalanus nasutus* (Goetze, 2003), which has a sister group along the North American coast, the rafting bryozoan *Membranipora* (Schwaninger, 2008), the giant kelp *Macrocystis* (Coyer et al., 2001) or, most evident, the bull kelp *Durvillaea antarctica* (Fraser et al., 2009). The latter shows distinct haplotypes along the central and northern Chilean coast. In analogy to results from New Zealand, they are most probably related to continuous warming of sea-water (Fraser et al. 2009, 2011). The observations on *D. antarctica* are consistent with further zoogeographic studies of littoral benthic and of pelagic organisms, which prove three distinct, most probably SST-related zoogeographical regions along the Chilean coast (Escribano et al., 2003; Hinojosa et al., 2006). Since goose barnacles from the southern hemisphere are abundantly rafting on detached *Macrocystis* and *D. antarctica* (Thiel & Gutow, 2005; Hinojosa et al. 2006), the coastal Chile subgroup appears to be very plausible.

In spite of extreme genetic divergence between populations of the bull kelp *Durvillaea antarctica* from Chile and New Zealand and further genetic differentiation within both regions, a genetically homogeneous population exists further south within the ACC (Fraser et al. 2009, 2011). Unified Antarctic genotypes were also described for the chaetognath *Eukrohnia hamata and the* Antarctic krill *Euphausia superba* (Bortolotto et al., 2011; Kulagin et al., 2014). Therefore, it might be hypothesized that a third, truly sub-Antarctic *L. australis* subtype might exist south of the Subtropical Convergence. However, one tiny *L. australis* specimen collected from floating *Durvillaea* kelp at around 50°S off the Chilean coast does neither belong to the postulated sub-Antarctic nor to the Coastal Chile subgroup, but it is more akin to the Southern Ocean subgroup.

### Lepas anatifera

Three major subgroups are discerned:

1. a subgroup collected off the Chilean coast during several sampling campaigns is set apart from all other globally sampled specimens by divergence in mitochondrial and nuclear markers, except for a single specimen. It shares the global nuclear genotype, but is retrieved inside the Chile group based on its mitochondrial genes. That specimen might be introduced by human activity;
2. a clear separation of *L. anatifera* into distinct Atlantic, Indo-Pacific, and NE Pacific sub-populations based on the mitochondrial loci within one nuclear genotype;
3. a subgroup containing specimens from dispersed global sampling sites. This biogeographic fragmentation on a population level needs discussion of Cenozoic plate tectonic evolution, Pleistocene climate development, and connected evolution of modern oceanic current systems.

### Group 1: The ‘coastal Chile subgroup’

The ‘coastal Chile subgroup’ is equivalent to our findings in *L. australis* (Fig. 6). Obviously, the cold water system of the Humboldt and the associated upwelling system along the Chilean margin prevent exchange between confined populations of certain organisms and cause a own Chilean phylogeographic province with strongly restricted permeability. Internally it is subdivided by increasing SST toward north (see above). Besides *L. australis* also *L. anatifera* is part of that phylogenetic province. The influence of the Humboldt Current is well expressed in the that warmer water species, which at the Chilean coast is not recorded south of 33S (Hinojosa et al., 2006).

**Figure 6:**
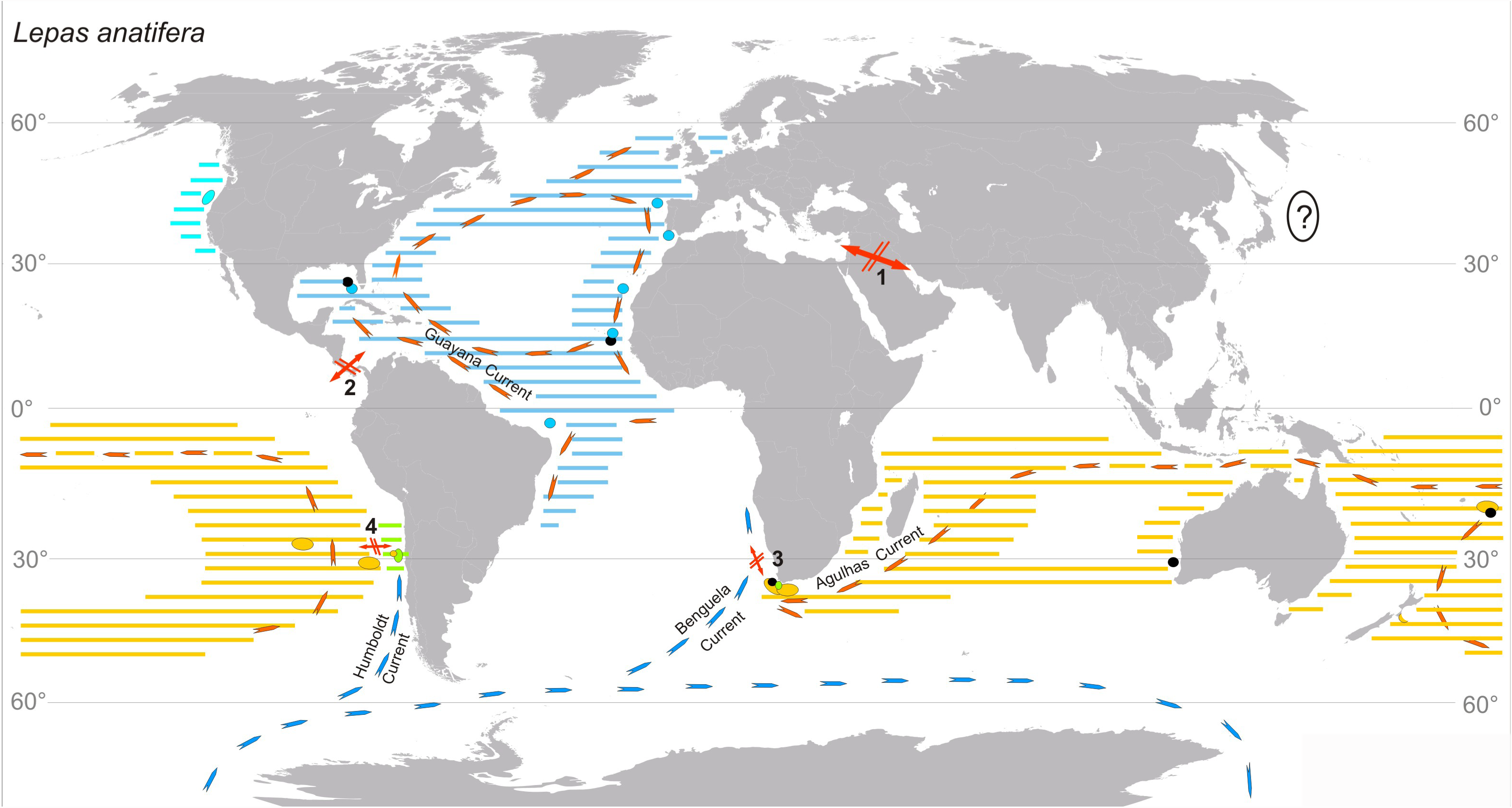
Sampling sites and retrieved population structure in *L. anatifera* in relation to our ocean current systems based on mitochondrial data. The only population set apart in the nuclear genotype is found at the Chilean coast (green).

The divergence of *L. australis* and *L. anatifera* in both mitochondrial and nuclear markers indicate an ancient origin of that ‘Chilean waterpocket’ which might be also seen in the tendency of floating debris to accumulate in the South Pacific west of Chile (Lebreton et al., 2012).

### Group 2 –Regional populations with mitochondrial differences

The Atlantic haplotype is a well individualized subgroup of *L. anatifera* (Fig. 6). It includes individuals from the Gulf of Mexico and specimens from the Spanish Atlantic coast (Galicia, Gulf of Cádiz), proving genetic exchange across the Atlantic via the Gulf Stream. Specimens further south from Madeira and from Cape Verde Islands demonstrate continuing rafting of that group within the Canary Current and thus well expressed circling within the North Atlantic gyre. Specimens from Brazil, however, also demonstrate trans-equatorial exchange with the southern hemisphere and thus a single large trans-equatorial Atlantic population. In the relatively narrow, north-south orientated Atlantic the North-Equatorial countercurrent is seasonally less pronounced and water masses of both hemispheres are well exchanged by the western, north-flowing Guiana Current in boreal winter and spring, though some throughflow occurs also in summer (Csanady, 1990; Condie, 1991). That current system is also in line with data on the distribution of *L. pectinata* in the Atlantic (see below). A similar pan-Atlantic population has also been observed in the ocean skater *Halobates micans* (Andersen et al., 2000) and in the scyphozoan *Pelagia noctiluca* (Miller et al., 2012).

The most favourable, ancient Tethyal gateway between Pacific and Atlantic marine populations was blocked by formation of the Panama land bridge about 3.5–3.1 million years ago (Myr) (Duque-Caro, 1990). Resulting divergence in marine taxa is known since decades and was updated and well elucidated by (Knowlton & Weigt, 1998) and recently by (Bacon et al., 2015). Due to SST as low as 8°C (Boyer et al., 2005) drifting around Cape Hoorn is most unlikely for the warmer water dwelling *L. anatifera*. Along the Chilean coast, barnacle population show *L. anatifera* dominating at 27S at SST of 21.21.9C, but fading towards 33S at SST of 18.21.0C (fig. 5 in Hinojosa et al., 2006). This southern limit is in accord with the GBIF database, which lists only a single specimen without collecting data further south at 43.88°S. Although somewhat varying, SST around the southern tip of South America have not been substantially higher through geological time (Mashiotta et al., 1999a; King & Howard, 2000; Feldberg & Mix, 2002; Droser et al., 2002) and therefore this passages apparently was continuously blocked.

The only feasible remaining genetic exchange between populations from the Indo-Pacific and Atlantic would be around the Cape of Good Hope. The warm Agulhas Current transports organisms from the Indian Ocean to the tip of South Africa, but is reflected back east there, thus impeding transport of rafting organisms into the Atlantic. Retroflection of the Agulhas Current is caused by West Wind drift and the Benguela Current, which transports cold water masses across the southern Atlantic towards the Southwest-African coast feeding the upwelling system along the southwest African (Namibian) coast (Lange et al., 1999). SST of the Benguela Current close to the coast are around 15°C (Clement & Gordon, 1995). Towards the open ocean, they rise up to 19– 20°C during summer, but do not exceed 17–18°C in winter (e.g. GES DISC 2012). These relatively cold waters might be outside of the preferential ecospace of *L. anatifera*, or warm-water seasons might be too short for successful rafting across the area. Besides current reflection these seem to be additional factors impeding gene flow. In fact, *L. anatifera* is reported in a diversified assemblage of goose barnacles from the Cape Town peninsula at the westernmost waning Agulhas Current (Whitehead & Biccard, 2011), but no record from the Namibian coast is known to us from literature or databases; only three specimens offshore Luanda (Angola) are listed in the GBIF database, the only ones along the African coast south of Dakar (Senegal) and south of our samples from approximately the same latitude from the Cape Verde Islands, which plot in Group 3 (see below). The formation of Indo-Pacific and Atlantic echinoid species within the genus *Tripneustes* was also related to the Benguela Current (Lessios et al., 2003). Direct comparison with *Lepas* might be somewhat problematic, since the pantropical shallow-water echinoids and their planktonic larvae are clearly more sensitive to cold water, but the current systems appears to be also an ancient and impermeable barrier to the ocean skater *Halobates micans* (Andersen et al., 2000) and to warm-temperate fish (Henriques et al., 2014). It also was discussed as a barrier for the nudibranch *Glaucus atlanticus* (Churchill et al., 2014). In fact, the Benguela Current System was established about 10–12 Myr ago (Diester-Haass et al., 2002; Heinrich et al., 2011). During the last glacial maximum the southern 15°C SST isotherm, herein considered as the minimum threshold temperature for *L. anatifera*, was considerably farther north than today, effectively touching the African and West Australian Coasts ({Williams:1998vx}; see our Fig. 1 for present day SST isotherms). Similar shifts have to be assumed for preceding Pleistocene glacials. These extended intervals of globally decreased sea surface temperatures provided first-order barriers to gene flow in *L. anatifera*; the ‘Algulhas Leakage’ during Pleistocene interglacials that allowed certain species to enter the Atlantic from the Indic (Vermeij, 2012) obvious did neither work for *L. anatifera* nor for *Glaucus atlanticus* (Churchill et al., 2014). In summary, the present-day Atlantic sub-population of the species (Brasil, Gulf of Mexico, Atlantic Spain, and Madeira) is related to different ancient vicariance events, namely (1) plate tectonics (closure of the Panama land bridge), (2) installation and modern persistence of current systems (Benguela Upwelling System), (3) climate variability in time resulting in shifting ecospace limits (extension of low SST towards the north during glacial periods) as well as to permant climate barriers (Cape Hoorn passage).

The Oregon coast subgroup of *L. anatifera* appears to be related to the North Pacific gyre (Fig. 6). It appears to be effectively separated by the Equatorial countercurrent from the Southern Pacific and Indic regions. Unfortunately, we do not have samples of the species off the Japanese islands. Therefore we cannot comment on relations between northwestern and northeastern Pacific. However, our findings seem to support the importance of the Pacific gyre systems for *L. anatifera* - just opposite than in *L. ansifera* (see below). Species of the ocean skater *Halobates* as well as genetically discriminated populations of the species *H. micans* and *H. sericeus*, (Andersen et al., 2000) (Leo et al., 2012), also prove disjunct northern and southern Pacific species, resp. populations. Data from the rafting bryozoan *Membranipora* shows a much higer degree of regionalization, but Northern and Southern Pacific clades are clearly separated, as well as a separate southeast Atlantic/south African clade (Schwaninger, 2008). Like *Membranipora*, also eucalanid copepods display strong regional diversification, based on mitochondrial and nuclear data (Goetze, 2003; 2005). *Rhincalanus nasutus* (Goetze, 2003) has five genetically distinct populations in the Pacific. Like for *L. anatifera*, a northeast Pacific population is discerned (our Oregon subgroup), but also a northwest Pacific population. In the southern Pacific, *Rhincalanus nasutus* shows separate southeast and southwest Pacific populations supplemented by a Sulu Sea population. Similarly *Eucalanus hyalinus* (Goetze, 2005) diverged into more regional populations than *L. anatifera.*

In fact, *L. anatifera* forms a common Indo-South Pacific subgroup. Our inclusion of South African individuals of *L. anatifera* demonstrate spatial extension of the occurrences into a huge Indo-Pacific population, similar to findings in the nudibranch *Glaucus. atlanticus* (Churchill et al., 2014). That realm represents the ‘Southern Hemisphere supergyre’ (Speich et al., 2007). Observational evidence of surface drifting tracer buoys also proved the existence of oceanic surface flow in the supergyre (van Sebille et al., 2012) and supports our observations.

These divergent patterns of *Lepas* and *Glaucus atlanticus* versus Halobates, *Membranipora* and the discussed eucalanid species demonstrate that the oceans have to be seen as highly dynamic and fragmented habitats for different species.

Group 3 - Specimens from dispersed global sampling sites

### Group 3: The dispersed ‘global’ group

Most puzzling is a globally distributed subgroup with specimens collected from the Gulf of Mexico, the Cape Verde Islands, South Africa (Cape Town), Western Australia, and Tonga. One could speculate that this is the most ancestral group originating from the circum-equatorial Tethys before disruption of this ocean by Miocene plate tectonic collisions resulting in closure of the Neotethys in the Middle East, about 10 Ma ago (e.g. (Mashiotta et al., 1999b)) and later closure of the Isthmus of Panama in the early Pliocene, about 3.5-3.1 Ma ago (Duque-Caro, 1990), vicariance effects also discussed for other taxa, like the ocean skater Halobates micans (Andersen et al., 2000) and the green alga Halimeda (Hillis, 2001). However, our results from the fast evolving mitochondrial genes is strongly contradictory. The specimens share the global nuclear genotype and thus we argue in favour of sampling artefacts introduced by anthropogenic or natural causes. Adults might have been transported attached to ships, or marine mammals, while larvae could be transported over long distance in ballast water or occasionally by sea birds. These are dispersal modes elucidated for more than half a century (e.g. (BISHOP, 1951)), but apparently they are still overridden by natural provincialism.

### Lepas testudinata

In general, *L. testudinata* is a rare species. From our nuclear data (18S gene) we retrieved *L. testudinata* as an ingroup to *L. anatifera*. This is contradicted by (Rees et al., 2014), who used 28S sequences in an assay to infer general barnacle phylogeny and found the species akin to *L. australis.* Given the morphological similarity of all three species we assume that they are very close and single gene genealogies cannot resolve their relationships and thus a phylogenomic study should be initiated in the future. Nevertheless, in *L. testudinata* the mitochondrial data suggests the formation two sub-groups, in South African and Australian waters, which is in contrast to *L. anatifera* and *L. australis*, where these regions are united. As discussed above, the connection between regions in *L. australis* will be mediated by the Antarctic current system and northeast deviating offsprings, where *L. testudinata* might not survive; ecological differences between *L. anatifera* and *L. testudinata* must remain the subject of further studies.

### *Lepas anserifera* and *L. pectinata*

Our 18S data place *L. anserifera* as an outgroup to *L. anatifera* and *L. australis,* and *L. pectinata* as an outgroup to all three (Fig. 3). Therefore, it is likely that – opposed to (Rees et al., 2014) – *L. anserifera* and *L. pectinata* are sister species. This is partly supported by morphology, especially by elevated ribs on shells of both species. Consequently, *L. anserifera* should share the tolerance of cooler waters found in *L. pectinata* by (Zevina & Memmi, 1981). This means that transit around South Africa and further through the Benguela upwelling system along the Namibian coast should be possible for both species, while the plate tectonic-induced barrier at the Isthmus of Panama and the climate-induced barrier at Cape Hoorn persist. The postulated similar ecological constrains explain the similar biogeographic signal for both species, which point to more or less undifferentiated global populations.

Concerning *L. pectinata*, our data contradict a split of into two antitropical populations as depicted by ((Hinojosa et al., 2006), fig. 6; see our Fig. 2) but support previous observations from the Atlantic, where the species occurs in temperate and tropical waters from the North of Ireland to the Cape Hoorn (Gruvel, 1905), (Young, 1990); see also data from the GBIF database). Our coI data could suggest a split into subgroups in then Indo-Pacific and Atlantic. However, this is neither supported by the 16S nor the nuclear 18S genotype and therefore we assume a global panmictic population of the species.

Also in *L. anserifera* a panmictic population on both hemisphere is documented by samples from the north-equatorial Atlantic (Cabo Verde, Senegal), from South Africa and Japan. Concerning the Indopacific realm, the difference to *L. anatifera* is striking, The latter, like other pelagic invertebrates discussed above, is strongly related to the hemispherical gyre systems, which are effectively separated by the eastward directed Equatorial Counter Current. However, in a study using tracer buoys and subsequent modelling the development of the huge garbage patches that accumulate in the gyre systems of the oceans on centennial timescales, (van Sebille et al., 2012) demonstrated the leakiness of that systems and interchange between all oceans. Most important appear to be non-linear mesoscale eddies (Chelton et al., 2011).

In the Pacific, trans-equatorial relations are also well-known for marine organisms. In a study on the calcareous sponge *Leucetta* ‘*chagosensis*’ Wörheide et al. (2002) found a trans-equatorial clade including individuals from the northern and central Great Barrier Reef as well as from Guam and Taiwan, though not further elucidated. Also (Benzie & Williams, 1997) found strongly related haplotypes of the giant clam *Tridacna maxima* from the Philippines and the Great Barrier Reef. Finally, (Williams & Benzie, 1998) noted remarkable analogies in the starfish *Linckia laevigata* between several southwest Pacific, Philippine and Japanese occurrences. Trans-equatorial dispersal was also mentioned by (Schwaninger, 2008) concerning Neogene dispersal of *Membranipora*, rafting on kelp southwards along the western coasts of the Americas and later northward in the Atlantic - a postulate opposite to the current system stressed by (van Sebille et al., 2012). Thus, additional pathways seem to exist/seem to have existed in the Pacific, seasonally, episodically or during times of changed climate or sealevel in the geological past. It might be speculated that during Pleistocene glacial periods and strongly restricted Indonesian throughflow toward the Indian Ocean a western transhemispherical current was developed, analogous to the present day Guiana Current along the western margin of the Atlantic.

In fact, Benzie & Williams (1997) and Williams & Benzie (1998) showed that the routes of gene flow in the western Pacific are not consistent with present-day ocean currents. They inferred dispersal during lowered Pleistocene sea levels and colder climate, which affected wind and current systems in multiple manners and general extension of species originating in the western Pacific in northwestern direction towards Southeast Asia.

## Conclusions

In spite of increasing molecular data on pelagic metazoans - true plankters or rafters - the emerging picture of biological partition of the oceanic water-masses and controlling factors remain contradictory and show different behaviour even for species within a single genus. The emerging picture is that the planktonic environment is a complex and diversified ecosystem. Truly cosmopolitan species seem to be extremely rare in the pelagic environment.

In this study we tried to obtain a global picture for the (epi-)pelagic metazoan genus *Lepas*, a supposedly cosmopolitan ecological generalist, which abounds in all major oceans. Detailed analyses of nuclear and mitochondrial sequence data in five species (*L. australis*, *L. anatifera*, *L. anserifera*, *L. pectinata*, *L. testudinata*) show a wealth of inherited geological factors and modern traits, which - in part interacting - result in the present distribution of the encountered *Lepas* populations. Our data indicate cryptic speciations in *L. australis* and *L. anatifera*, but at the same time global *L. pectinata* and *L. anserifera* and a potentially (if not anthropogenic) global *L. anatifera* population.

In *Lepas*, the most ancient split into different populations resulted from the closure of the Isthmus of Panama separating Pacific and Atlantic populations in *Lepas anatifera*. An still older vicariance event, the closure of the Tethys seaway by the plate tectonic collisions in the Middle East fragmenting a global stem group(?) of *L. anatifera* remains speculative. Pleistocene cold climates and sealevel lowstands in the insular world of the western Pacific resulted in different wind and current patterns. They induced migration patterns differing from present-day oceanic currents and explain transequatorial populations between the South Pacific and Japan in *L. anatifera*. Alternatively, or concomitantly, leakage of the South Pacific gyre along the western coast of South America has to be considered. Installation of the circum-Antarctic current system and especially of the Benguela Current in the later Miocene, around 10-12 Mio a ago, started to inhibit rafting of the species around the southern tips of South America and South Africa. Cold Pleistocene climates and extension of coldwater masses towards lower latitudes definitely closed these gateways for *L. anatifera*, which prefers warmer waters. Both gateways are still closed for the species in modern time. However, as seen in the common global gene pool, these temperature barriers do not affect *L. ansifera* and *L. pectinata*. Present-day current systems are responsible for further partioning of water masses. We stress the fact, however, that all current systems have extended geological histories, as herein amply discussed for the Benguela Current, and also seen in the genetically ancient L. australis and L. anatifera of the ‘Chilean waterpocket’, which is related to the Humboldt Current. The Pacific gyres are first-order entrapment systems. The North Pacific gyre results in separation of the Oregon population, though missing of East Asiatic (“Japanese”) individuals remain enigmatic. The South Pacific gyre appears to be strongly temparature controlled in its south-directed and north-directed segments, as seen in separate Chilean and New Zealand populations in *L. australis* as well as in a Chilean population of *L. anatifera* set apart from a general Indopacific population. Finally, the width of oceans and contiguous land-masses might favour trans-equatorial and transcontinental currents like the Guayana Boundary Current and the Gulf Stream, resulting in a comon *L. anatifera* subgroup in both hemispheres in the Atlantic. Last not least, anthropogenic transport and swimming mammals further complicate the distributional pattern.

Our results show that a multitude of inherited and persistent barriers exist in the present-day open ocean. They act on species level according to the ecospace realized by a taxon in the geological past and today. That implies the following consequences:

1. Speciation in the pelagic realm has its roots in geological time (see for example also Andersen et al., 2000; Fraser et al., 2009; Norris & Hull, 2011; Leo et al., 2012). The postulate of rapid diversification and regionalisation of zooplanktic species (Peijnenburg & Goetze, 2013) cannot be regarded as general valid rule.
2. Allopatric speciation is of prime importance, even in apparently barrier-less oceanic regions like in the southern Pacific gyre. Sympatry remains an option in the vast pelagic realms, but has to be proved.
3. There is no general pattern for cryptic speciation, resp. formation of populations, in the pelagic realm. Studies on species level are necessary for different taxa to counterbalance ecological limits and the corresponding barriers operating in time and space.
4. The existence of truly cosmopolitan taxa in the pelagic environment is not yet settled. Only very few taxa, such as *L. pectinata* and *L. ansifera*, which are, first, extreme ecological generalists and, second, capable of long-distance dispersal might be able to maintain gene flow throughout the worlds oceans. Both species might be model taxa to clarify that problem by extending regional data and 2nd generation sequencing approaches.

In fact, Darwin (1851; 1854) started the discipline of phylogeography. He was the first to note the morphological plasticity within species - notably within *Lepas -* and the importance of (land-)barriers for species separation. Some 120 years later, Newman (1972) recognized the regional diversification, respectively the formation of subspecies within *L. anatifera* based on morphological differences. His results are partly supported by our genetic data. Still, our state of the art on *Lepas* and the discussed comparisons with other taxa illustrate that the biotic dynamics in the world oceans are far from being understood. Being the largest biotope of the world and probably the most valuable resource for mankind, however, such data is indispensable, not the least concerning questions of conservation in times of global change (Wright et al., 2015).

## Acknowledgements

First results on the phylogeography of *Lepas* were obtained during work on the unpublished diploma thesis of PS (2009), but results presented herein were achieved in the course of other projects by low-cost research neither applied nor financed from research foundations. However, the authors acknowledge continued financial and laboratory support from the Volkswagen Stiftung (PS) and Universität zu Köln (PS, HGH). We are especially thankful towards Martin Thiel (Coquimbo, Chile) for providing specimens and valuable comments on the manuscript. We thank our student assistant A. Kraemer-Eis for help with PCRs and sequencing as well as figure design. We are grateful to all colleagues providing samples: P. Wirtz, Madeira; A. Biccard and C. Griffith, Cape Town; A. Hosie, Wellington and Perth (at the Western Australian Museum and acquired with help of Prof. Dr. M. Türkay, Frankfurt); as well as all others who we forgot to name here.

